# Mutafy: A webserver to identify high quality mutant protein structures in the Protein Data Bank

**DOI:** 10.1101/2023.03.22.533870

**Authors:** Deborah Ness, Jiajing Hu, Munishikha Kalia, Richard JB Dobson, Ammar Al-Chalabi, Alfredo Iacoangeli

**Affiliations:** Maurice Wohl Clinical Neuroscience Institute, Department of Basic and Clinical Neuroscience, King’s College London, London, UK; Department of Biostatistics and Health Informatics, King’s College London, London, UK; National Institute for Health Research Biomedical Research Centre and Dementia Unit at South London and Maudsley NHS Foundation Trust and King’s College London, London, UK; Institute of Health Informatics, University College London, London, UK

## Abstract

Changes in the amino acid sequence of proteins resulting from nonsynonymous variants in the genome, can have significant effects on protein folding, stability, dynamics, and function, which may ultimately lead to diseases. The analysis of large sets of disease associated variants is a common approach for the study of pathogenic mechanisms. *In-silico* mutagenesis experiments based on wildtype structures of target proteins are a common approach to this aim, however these do not account for the effect of variants on folding and might not accurately reflect conformational changes. A growing number of experimentally solved protein structures harbouring disease-associated mutations, including single amino acid variants, are deposited in the worldwide Protein Data Bank (PDB). Nevertheless, identifying high-quality structures for specific missense variants of interest remains challenging due to the growing number of deposited protein structures in the PDB, and the lack of a dedicated interface and annotation system to search and retrieve mutant protein structures. As a result, mutant protein structures in the PDB are a powerful source of information which is largely underused. To address these shortcomings, we have developed Mutafy, a publicly available webserver to identify high quality mutant protein structures. Given input human genes, the webserver finds structures of the corresponding coded wildtype proteins and their available solved mutants, selects high quality structures, annotates them with information from biomedical databases to favour their interpretation and selection, and allows for the interactive exploration of the results and 3D visualisation. Mutafy is publicly available without requiring user registration at https://mutafy.rosalind.kcl.ac.uk.

## Introductions

Genomic mutations resulting in changes in the amino acid sequence of the corresponding coded proteins can alter protein function, folding, stability and protein-protein interactions (1). Such effects can result in pathogenic mechanisms disrupting fundamental processes and lead to a variety of diseases (2). With the advancement of high-throughput technologies, genetic sequencing data are widely available in the biomedical field and as a result, the discovery of genetic mutations is often the starting point for their study (3,4). Understanding how amino acid changes affect the protein can provide insight into associated biological processes. Computational approaches based on force fields or biological annotations are widely adopted for this aim (5,6). For example, molecular dynamics is a common technique utilised to study how variants affect protein structures and their dynamic behaviour. These methods rely on the availability of high-quality structures of the wild-type (WT) and its mutants. The latter are usually available via *in-silico* mutagenesis approaches that generate mutant proteins by computationally changing the side chains of target amino acids (7). However, *in-silico* generated mutations do not account for the effect of mutations on folding and might not accurately reflect the conformational and functional changes caused by the mutations (8,9). This greatly limits the validity and generalizability of the results from studies of this type. The Protein Data Bank (PDB) is a publicly available database where an increasingly high number 3D structures of proteins, normally solved using X-ray crystallography or nuclear magnetic resonance (NMR), are deposited and made available to the research community (10). At present approximately 200,000 experimentally solved protein structures, many of which harbour mutations, are available in the PDB. Such “mutant” protein structures represent a great resource for the study of the impact of amino acid changes on proteins and their related biological processes. However, identifying all available mutants in the PDB would require a complex and time-consuming manual review of large sets of models given the high number of deposited structures and the lack of a dedicated interface and annotation system to search and retrieve them. Previous efforts to make mutant proteins, whose structures were solved “*in-situ*”, easily accessible were limited. To our knowledge current databases of protein mutations focus on the identification of protein sequences and are usually focused on specific domains, e.g. cancer variants (11). The Protein Mutant Database (PMD) (12) was the only viewing and retrieving system with a broader scope, however, it focused on the annotation of literature and identification of mutant sequences from published articles that were subsequentially linked to PDB models which did not necessarily harbour the mutations. Despite gaining substation popularity after its release in 1999, it has been discontinued for years.

In order to facilitate the identification and selection of experimentally solved structures of mutant proteins, we have developed Mutafy, a user-friendly webserver for on-the-fly identification of high-quality mutant protein structures in the PDB. Mutafy takes as input a list of HUGO Gene Nomenclature Committee (HGNC) gene symbols (13) and screens the whole PDB for all structures associated with those genes, identifying and annotating them with any amino acid changes with respect to the reference sequences from Uniprot (14). The results are made available via interactive tables and a protein visualization tool (15) on the webserver. Structures for all available mutations and the wildtype are ranked and selected according to their resolution. Alphafold (16) predicted models of the WT are also retrieved. All structures can be downloaded from the webserver and summary results tables are emailed to the user. Mutafy is publicly available without requiring user registration at https://mutafy.rosalind.kcl.ac.uk.

## Results

### Pipeline overview

Mutafy (Figure 1) takes a list of gene names (HGNC gene symbols) as input and identifies all associated human protein structures available in the PDB. The sequences of these PDB entries are then aligned with BLASTp (17) against the canonical protein reference sequence (obtained from UniProt) to identify the correct proteins and to remove those which do not pass the WT similarity thresholds (minimum proportion of residues that align to the reference sequence and minimum length of the protein sequence), and to identify mutations. Mutations are classified as single amino variants (SAVs) if they involve only one amino acid. The identified mutations are annotated using ClinVar (18) to highlight their clinical relevance and interpretation. Other pieces of information about the model such as percentage of solved residues, resolution, and experimental method for the structure determination, are extracted from the PDB. This information is made available to the user, and it is used for the selection of models presented on the webserver. Mutafy generates 4 distinct output tables listing the protein structures with the highest resolution available for the input genes, and grouping structures according to their mutations in the following way:

**Figure 1.**
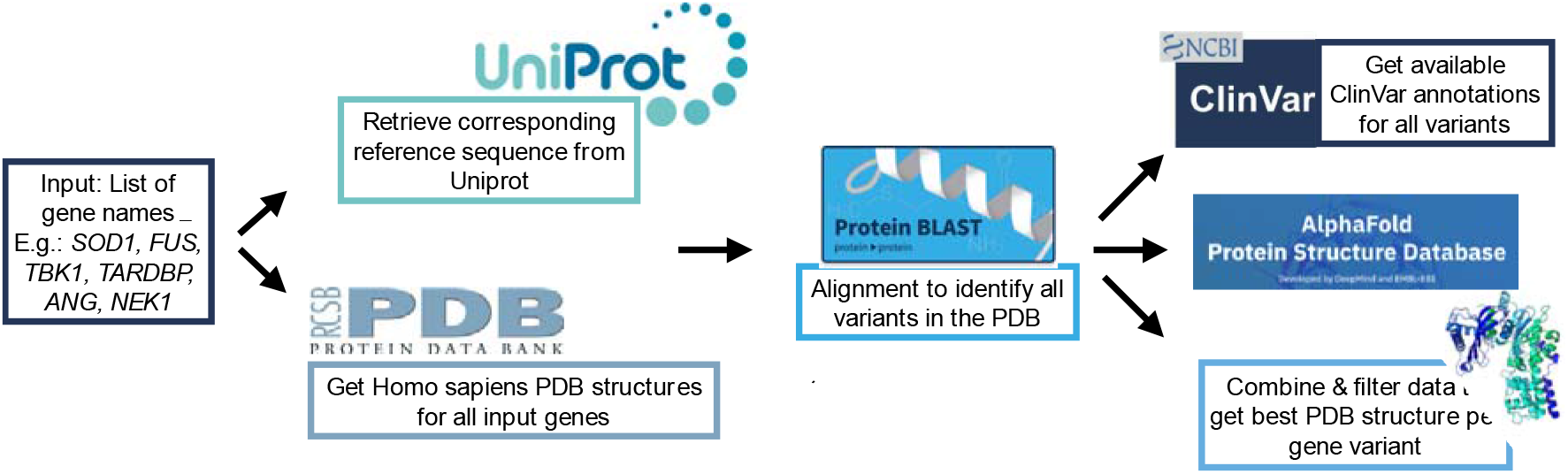
Flowchart of the pipeline run by Mutafy

1. Best structure per single amino acid variant (SAV): the structure with the highest resolution of each single amino acid variant (SAV) for all proteins associated with the input genes, i.e. structures which have only one amino acid substitution in their sequence compared to the WT protein sequence.

2. Best structure per unique sequence / combination of mutations: the best available structure for each unique sequence in the PDB, including WT, associated with the input genes.

3. Best structure per any identified mutation (regardless of other mutations coexist in the same protein): the best available structure for each mutated residue found in any of the PDB structures. The same structure might be selected for more than one amino acid change if their sequence harbours 2+ mutations.

4. Wildtype structure: all available WT structures associated with the input genes and ranked according to their resolution.

### Webserver overview

#### Homepage

Mutafy’s homepage is minimal and intuitive (Figure 2A). It presents a brief explanation of the pipeline for the identification of the models, an input text field for the user to fill with the list of target genes, and another for the optional email address to notify when the job is complete and to send the summary results. A simple job can be run by simply typing a gene name and clicking on the “Submit” button. An example gene list can be run by clicking on the “Example” button and gene names are automatically suggested when the user starts typing in the corresponding input text field. Advanced options are available in the tick-to-show hidden menu for expert users. Help messages with brief descriptions of the advanced options are also available. The tutorial page provides a step-by-step guide for the use of Mutafy and a detailed description of its components.

**Figure 2.**
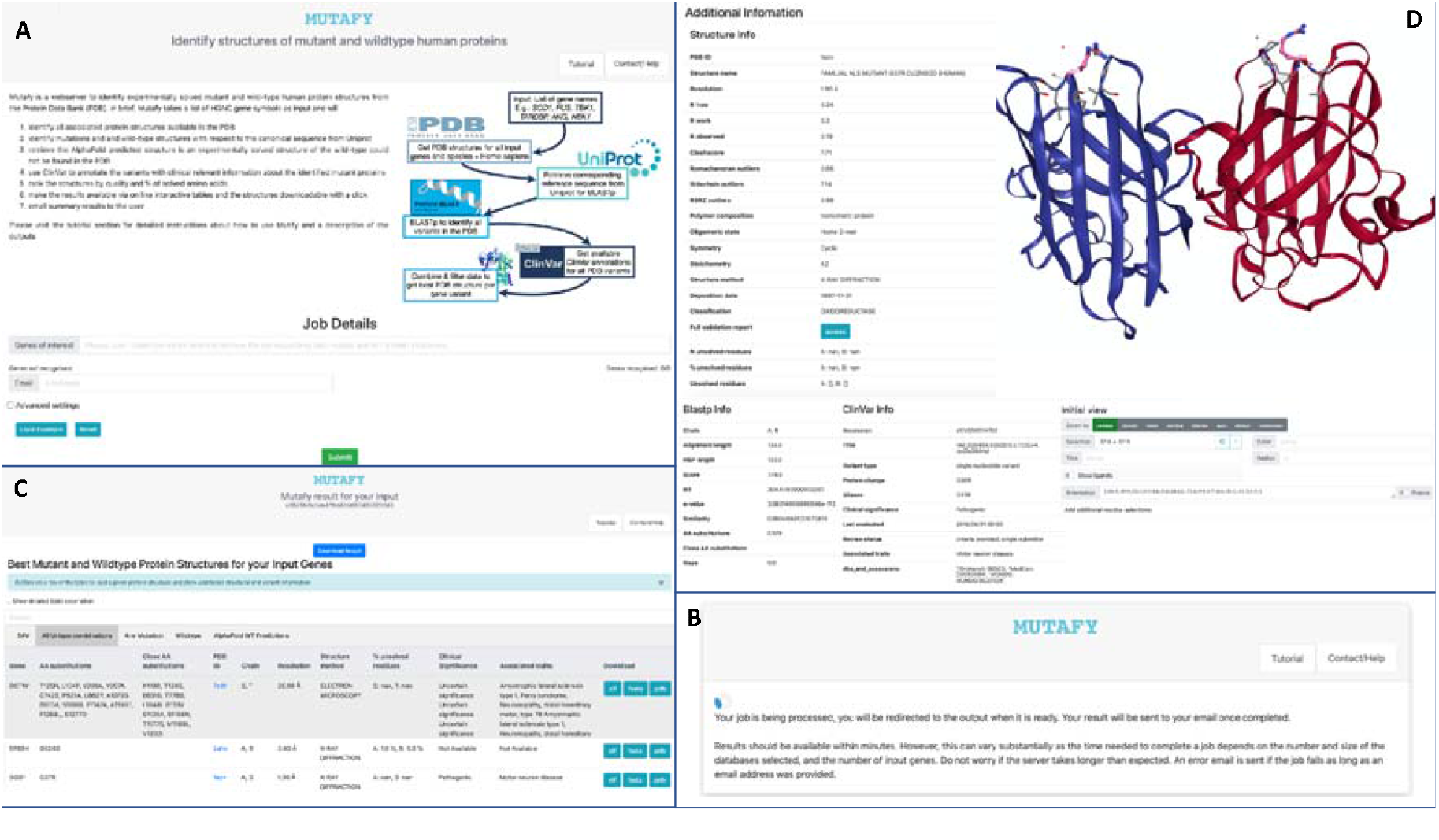
Overview of the webserver: A) Homepage. B) After submission waiting page. C) Results page; example of interactive results table. D) Results page; example of interactive protein visualisation. By default, the side chains of the mutation are highlighted.

#### After submission

After submission, the user is directed onto the waiting page (Figure 2B) where the unique job ID is displayed. The ID can be used in the homepage to retrieve the job results. If a valid email address was provided at submission, the job ID and a link to the results page are also emailed. The waiting page automatically refreshes every 10 seconds until the job is completed. The user is then redirected to the results page. A standard job takes about 10 minutes to be completed. However, jobs can take longer as the processing time depends on the number of genes and identified PDB structures that need to be parsed. However, users are notified via email when a job is completed. Results are saved in the server database so that Mutafy can re-use the results from previous jobs if the same genes were submitted with the same options and no related new entries are found in the PDB. Submissions of genes present in the server database normally complete within minutes.

#### Results page

The results page consists of five tabs (Figure 2C). Four of them interactively display simplified versions of the output tables that are emailed to the user and were described in the “Pipeline overview” section. The fifth table lists the Alphafold models of the WT. For each structure, gene name, amino acid change, list of chains in the pdb file, resolution, structure determination method, percentage of unsolved residues and ClinVar annotations are provided. Additionally, direct links to the corresponding PDB page, and direct download buttons for the structure (PDB), sequence (fasta) and crystallographic information file (cif) are also provided. By clicking on any structure of the lists, the server initialises an interactive visualisation tool that allows for the exploration of the structure and its variants, as well as the full set of information fields from the BLAST alignment to the reference, ClinVar annotation and the PDB entry (Figure 2D).

### Analysis of ClinVar genes

ClinVar is an established and widely-used database that links information about genomic variants to their clinical significance, i.e. whether variants are considered pathogenic, likely pathogenic, of uncertain significance, likely benign or benign, and the related supporting evidence for such interpretations. ClinVar often represents the starting point for the selection variants in mutagenesis studies aimed at investigating the molecular basis of human diseases. We have analysed all ClinVar genes to test the availability of clinically relevant mutant protein structures in the PDB and to provide the webserver database with a substantial number of pre-processed entries of wide interest that can be used to reduce the time needed by jobs that include one or more such genes (Figure 3). 4751 genes were found in ClinVar and submitted to Mutafy. At least one related PDB entry was identified for 2709 genes. For these 2709 genes Mutafy retrieved 1660 mutant protein structures in the PDB and 10.91% of these harboured mutations with a reported clinical interpretation in ClinVar.

**Figure 3.**
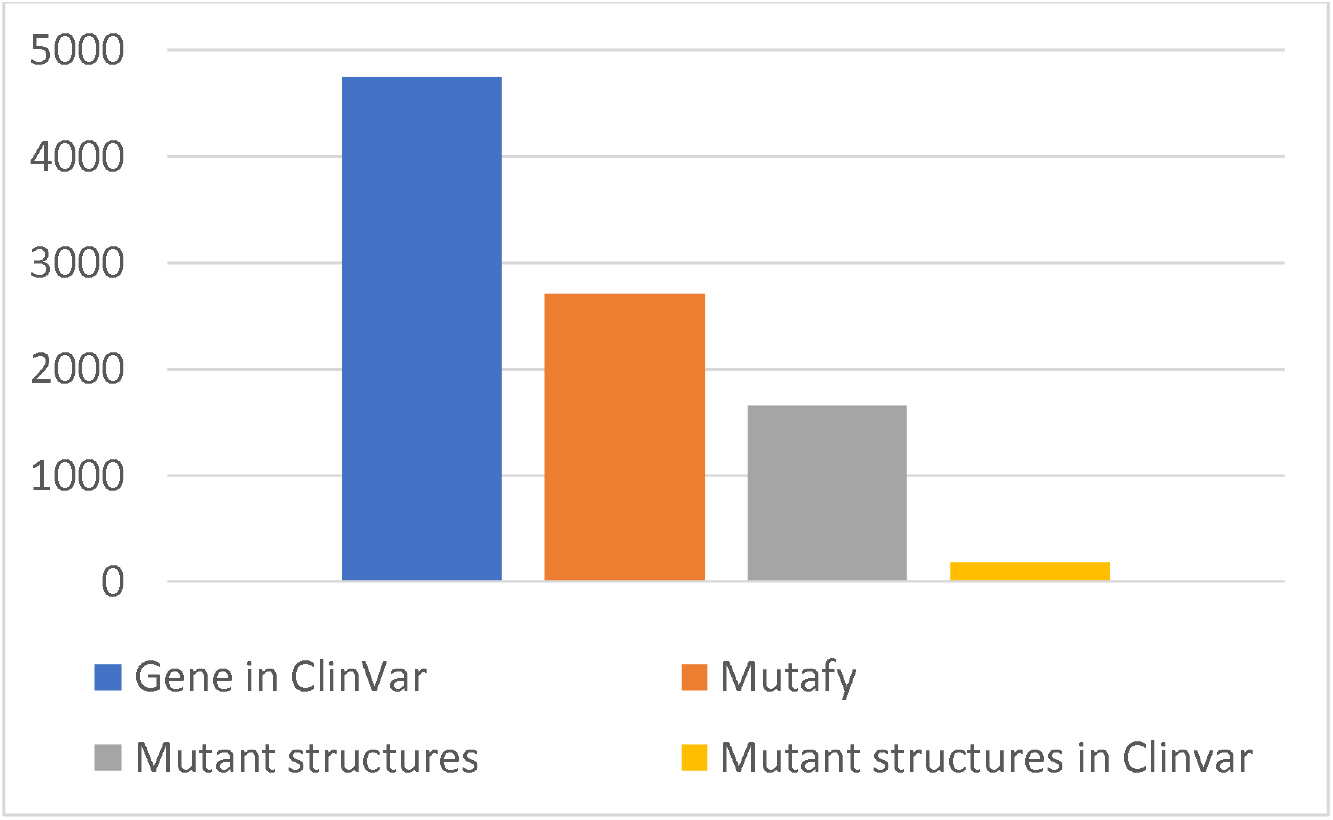
Overview of Mutafy analysis of CLinVar gene. Total number of genes in ClinVar (blue); number of ClinVar genes for which Mutafy could find and select at least one structure in the PDB (orange); Total number of mutant structures that Mutafy found in the PDB for the ClinVar genes (grey); mutant structures that harbour a variant present in ClinVar (yellow).

## Discussion

The generation of sets of experimentally solved protein structures harbouring mutations linked to biological processes such as human diseases, is a critical step for the design of studies which aim to investigate the underlying mechanisms. Mutafy streamlines this process, making it straightforward to perform without the need for informatics training and minimizing the time required. Its user-friendly interface allows users to submit jobs with a simple selection of the target genes and a click, and its results can be explored and visualised on the webserver via interactive tables and a protein 3D structure visualisation tool. We believe that Mutafy can be a valuable tool for a broad audience of biomedical scientists, and it fills an important gap in the landscape of current bioinformatics webservers.

### Webserver and Code availability

The Mutafy webserver is available at the following URL https://mutafy.rosalind.kcl.ac.uk/ The code and documentation for a local implementation of the pipeline are available on GitHub (19): https://github.com/Utilon/MutaPipe_Repo. The webserver is based on Michelanglo (20) and its codebase is available on GitHub: https://github.com/CMD-Oxford/Michelanglo-and-Venus.

## Materials and Methods

### Webserver implementation

The webserver was developed starting from the codebase of Michelanglo, which is a Python3 webapp running the Pyramid web framework (21), and adapted to reflect the Mutafy input/output and graphical requirements.

### MutaPipe

Mutafy runs an adapted version of the MutaPipe pipeline to identify mutant and WT protein structures. Given a set of gene names provided as HGNC symbols, MutaPipe identifies all associated PDB IDs (exact search for species and gene name = Homo sapiens using the PDB API) and downloads the corresponding fasta, mmCif, and pdb files to extract and combine relevant information (incl. resolution, sequence, unsolved residues in PDB structures). Then the canonical protein sequences (stored in a fasta file downloaded via the UniProt FTP client) for all input genes with available PDB data are downloaded. BLASTp is used (e-value threshold: 0.05) to find mutations in the sequences of all identified PDB structures. Biopython (22) includes the Bio.Blast.Applications module which provides wrappers for the NCBI blast tools (23), and Mutafy uses the standard NCBI default settings (the same as on the NCBI website) to perform a BLASTp of two sequences at a time (one from a protein structure and one being the canonical protein sequence identified from UniProt).

BLASTp is performed separately for every sequence in a given structure (Note: PDB structures may consist of multiple chains, each with a different amino acid sequence) against the UniProt reference sequence for the input gene associated with this structure (e.g. BLASTp of all sequences in a given FUS structure against the canonical protein sequence for FUS). This procedure allows Mutafy to promptly discard as models all sequences within a structure which do not share significant similarity with the reference WT sequence. Mutafy has two adjustable options allowing users to define threshold values for this selection:

Minimum relative sequence length: Mutafy will compare the length of each protein sequence (or chain) in a protein structure (which has been associated with the input gene) to the length of the canonical protein sequence (canonical/WT protein sequence encoded by the input gene). The minimum relative sequence length defines the threshold sequence length compared to the WT/canonical sequence in percentage, under which proteins will not be considered adequate models anymore. Hence this feature allows users to exclude models from the output which have a shorter amino acid sequence than a given percentage of the reference sequence, e.g. the default value 0.5 results in all models listed in the output tables having a sequence which is at least 50% as long as the corresponding canonical sequence. Changing this value might be of interest if one is interested in retrieving shorter fractions of a large protein.

Minimum HSP coverage: A high-scoring segment pair (HSP) is a subsegment of a pair of sequences that share high levels of similarity, i.e. without many mismatches or gaps. A BLASTp will identify 0-n HSPs, depending on (a) the similarity of the two aligned sequences, and (b) the settings of the local or global alignment algorithm which generated them (NCBI default settings). This feature allows user to exclude as models all sequences whose best High Scoring Segment Pair (HSP) covers less than a given percentage of the reference sequence, e.g. if the value is set to 0.45, all protein sequences which do not have a subsegment with high similarity to the canonical WT sequence, covering at least 45% of the canonical sequence length, will not be considered adequate models and will be excluded from the output. Changing this value might be of interest when searching for shorter peptides or fractions of a larger protein too. BLASTp results are used to extract information (incl. e-value, bit, alignment length, mismatches, close mismatches, gaps, indels) needed by Mutafy to identify / rank all unique protein sequences in the PDB associated with the input genes. Moreover, Mutafy uses Entrez Direct to download additional information from the ClinVar database, including data on variant pathogenicity and associated traits, for all available variants for all input genes with available PDB data. Whenever a mutation resulting in an amino acid change is identified in a PDB structure, Mutafy will check if ClinVar stores corresponding information for this mutation and annotates variants wherever possible. Mutafy will order and rank the filtered and annotated structures primarily according to their resolution (and secondarily by additional quality parameters in the background, e.g., similarity) and put together multiple output tables containing the best structures for all mutant and WT sequences available for the input genes. In case the user would like to include more than just the top/best structure in the output, the option “Number of top structures” in the Advanced Settings sections on the Mutafy input page can be adjusted (default = 1). Mutafy generates 4 distinct output tables listing the highest quality (highest resolution) protein structures available for the input genes as described in the Pipeline overview section of the results.

## Acknowledgements

JU is funded by the China Scholarship Council (CSC) via the King’s-China Scholarship Council PhD Scholarship programme. RJBD is supported by the following: (1) NIHR Biomedical Research Centre at South London and Maudsley NHS Foundation Trust and King’s College London, London, UK; (2) Health Data Research UK, which is funded by the UK Medical Research Council, Engineering and Physical Sciences Research Council, Economic and Social Research Council, Department of Health and Social Care (England), Chief Scientist Office of the Scottish Government Health and Social Care Directorates, Health and Social Care Research and Development Division (Welsh Government), Public Health Agency (Northern Ireland), British Heart Foundation and Wellcome Trust; (3) The BigData@Heart Consortium, funded by the Innovative Medicines Initiative-2 Joint Undertaking under grant agreement No. 116074. This Joint Undertaking receives support from the European Union’s Horizon 2020 research and innovation programme and EFPIA; it is chaired by DE Grobbee and SD Anker, partnering with 20 academic and industry partners and ESC; (4) the National Institute for Health Research University College London Hospitals Biomedical Research Centre; (5) the National Institute for Health Research (NIHR) Biomedical Research Centre at South London and Maudsley NHS Foundation Trust and King’s College London; (6) the UK Research and Innovation London Medical Imaging & Artificial Intelligence Centre for Value Based Healthcare; (7) the National Institute for Health Research (NIHR) Applied Research Collaboration South London (NIHR ARC South London) at King’s College Hospital NHS Foundation Trust.

AAC is an NIHR Senior Investigator (NIHR202421). This research is part an EU Joint Programme - Neurodegenerative Disease Research (JPND) project. The project is supported through the following funding organisations under the aegis of JPND: Medical Research Council; Economic and Social Research Council and the Motor Neurone Disease Association. Funding for open access charge: UKRI.

A.I is funded by the Motor Neurone Disease Association, South London and Maudsley NHS Foundation Trust, MND Scotland, National Institute for Health Research, Spastic Paraplegia Foundation, Alzheimer’s Research UK, Darby Rimmer MND Foundation and Rosetrees Trust. This study represents independent research partly funded by the National Institute for Health Research (NIHR) Biomedical Research Centre at South London and Maudsley NHS Foundation Trust and King’s College London. We acknowledge use of the research computing facility at King’s College London, Rosalind (https://rosalind.kcl.ac.uk) and CREATE, which are delivered in partnership with the National Institute for Health Research (NIHR) Biomedical Research Centres at South London & Maudsley and Guy’s & St. Thomas’ NHS Foundation Trusts and part-funded by capital equipment grants from the Maudsley Charity (award 980) and Guy’s and St Thomas’ Charity (TR130505). The views expressed are those of the author(s) and not necessarily those of the NHS, the NIHR, King’s College London, or the Department of Health and Social Care.

## Notes

### Competing Interest Statement

The authors have declared no competing interest.

